# Maternal hyperglycemia impairs postnatal skeletal muscle growth and induces stage-dependent gene expression changes in mouse offspring

**DOI:** 10.64898/2026.10.03.755574

**Authors:** Yayoi Tokunaga, Tamaki Kurosawa, Taiki Mihara, Delphine Duprez, Masatoshi Hori

## Abstract

Maternal hyperglycemia can affect offspring development and metabolism, but how its effects on skeletal muscle change during postnatal life remains unclear. We investigated skeletal muscle phenotypes in male offspring of streptozotocin-induced hyperglycemic mice at 4 and 8 weeks of age, representing early post-weaning and later postnatal stages, respectively. Maternal hyperglycemia resulted in lower offspring body and skeletal muscle weights at both time points. Myofiber cross-sectional area was reduced without a reduction in total myofiber number, indicating impaired myofiber growth rather than myofiber loss. At 4 weeks, skeletal muscle showed type I/IIA hybrid myofibers and increased expression of fetal, myogenic, and proliferation-associated genes. This immature profile was not maintained at 8 weeks; instead, the transcriptional profile shifted toward reduced mitochondrial and oxidative metabolism. DNA methylation changes at 4 weeks showed limited correspondence with transcriptional alterations. Taken together, maternal hyperglycemia impaired postnatal myofiber growth at both time points, with distinct molecular profiles at 4 and 8 weeks.

## Introduction

Maternal hyperglycemia during pregnancy can influence fetal development and produce lasting effects on offspring growth and metabolism [1]. In humans, gestational diabetes mellitus is commonly associated with maternal insulin resistance and increased fetal growth [2]. In contrast, severe experimental maternal diabetes can result in fetal growth restriction, pregnancy loss, and reduced postnatal growth [3–5].

STZ-treated rodents are commonly used to examine the effects of maternal hyperglycemia on offspring development. STZ produces insulin-deficient maternal hyperglycemia rather than the obesity-associated insulin resistance that typically underlies human gestational diabetes mellitus. Moreover, the maternal and offspring phenotypes produced by STZ vary with the timing and severity of diabetes. Severe STZ-induced maternal diabetes has been associated with reduced litter size and impaired fetal growth [3, 4, 6], whereas milder experimental hyperglycemia can produce different growth and metabolic outcomes. Thus, STZ-induced maternal diabetes should not be regarded as a direct surrogate for human gestational diabetes but provides an experimental model for examining the consequences of a defined maternal hyperglycemic environment on offspring development.

Previous studies using STZ-induced maternal diabetes have demonstrated persistent metabolic abnormalities in offspring, including altered glucose homeostasis, insulin responsiveness, and peripheral glucose utilization [7, 8]. Skeletal muscle is particularly relevant to these phenotypes because it is a major site of insulin-stimulated glucose disposal and plays a central role in whole-body glucose homeostasis [9]. Altered skeletal muscle insulin signaling and glucose metabolism have been reported in offspring exposed to maternal diabetes or hyperglycemia [10, 11]. However, skeletal muscle is not only a metabolic organ but also undergoes extensive myofiber growth and maturation during the early postnatal and prepubertal periods [12]. Whether severe maternal hyperglycemia affects this postnatal growth and maturation process, and whether the associated molecular alterations persist or change across postnatal development, remain incompletely understood.

Changes in postnatal muscle phenotype may be accompanied by persistent or stage-dependent alterations in gene regulation. Developmental exposure to an altered maternal metabolic environment can induce epigenetic modifications, including DNA methylation [13–15]. DNA methylation could therefore contribute to transcriptional alterations observed in offspring skeletal muscle, although the extent to which differential gene expression after maternal hyperglycemia corresponds to concurrent DNA methylation has not been well characterized.

In the present study, we used an STZ-induced severe maternal hyperglycemia model to characterize postnatal metabolic and skeletal muscle phenotypes in male offspring. Offspring were examined at 4 and 8 weeks of age to determine whether impaired skeletal muscle growth persisted across postnatal development and whether the associated molecular phenotype differed between these stages. We assessed systemic glucose-related phenotypes, skeletal muscle growth, myofiber morphology, and fiber-type composition and performed transcriptomic analyses at both time points. Because extensive transcriptional alterations were observed at 4 weeks, we further examined their relationship with concurrent DNA methylation changes using reduced representation bisulfite sequencing.

## Methods

### Animals

All animal experiments were approved by the Animal Care and Use Committee of the Graduate School of Agricultural and Life Sciences, The University of Tokyo (approval no. P-21-107), and were performed in accordance with institutional guidelines for animal experimentation. Male and female C57BL/6J mice aged 8–12 weeks were purchased from Shiraishi Laboratory Animals Co., Ltd. Mice were housed at 2–4 animals per 330 cm² cage under controlled conditions at 22 ± 2°C with a 12-h light/dark cycle and had ad libitum access to food and water.

### Streptozotocin-induced maternal hyperglycemia model

Two female mice and one male mouse were housed together at 17:00, and mating was assessed the following morning at 09:00. Pregnancy day 0 was defined as the day on which sperm were detected by vaginal smear examination. To induce maternal hyperglycemia, streptozotocin (STZ; FUJIFILM Wako Pure Chemical Corporation, Japan) was dissolved in ice-cold 0.1 M citrate buffer and administered intraperitoneally at 100 mg/kg once daily for 3 consecutive days. Control mice received an equivalent volume of citrate buffer. Blood glucose levels were measured using a glucose meter (LAB Gluco, Research and Innovation Japan Co., Ltd., Japan) with blood collected from the tail vein. Mice with blood glucose levels >250 mg/dL between 5 days after STZ administration and gestational day 15 were classified as hyperglycemic and included in the study. Mice that did not develop hyperglycemia were excluded. Maternal health status, activity, coat condition, and access to food and water were monitored approximately three times per week, while blood glucose and body weight were monitored weekly. Mice showing >20% body-weight loss 7 days after STZ administration were euthanized under deep isoflurane anesthesia by cervical dislocation according to the predefined humane endpoint. Male offspring born to vehicle-treated dams were designated as the control group, whereas male offspring born to hyperglycemic dams were designated as diabetes mellitus offspring (DMO). Offspring were analyzed at 4 and 8 weeks of age.

### Sample collection

At 4 and 8 weeks of age, male offspring were euthanized by cervical dislocation under deep isoflurane anesthesia. Blood, pancreas, liver, and hindlimb skeletal muscles were collected. The tibialis anterior (TA), gastrocnemius, and soleus muscles were carefully dissected and weighed. The combined length of the femur and tibia was measured as an index of longitudinal skeletal growth. Pancreatic tissues were processed for histological analysis as described below. TA muscles used for histological analysis were mounted on cork with tragacanth gum and rapidly frozen in isopentane precooled in liquid nitrogen. Liver and skeletal muscle samples used for RNA analysis were immediately frozen and stored at −80°C until RNA extraction.

### Serum preparation and insulin measurement

Blood samples were incubated at room temperature for 2 h and centrifuged at 2,000 × g for 15 min at 4°C. The resulting supernatant was collected as serum. Serum insulin concentrations were measured using a Mouse Insulin ELISA Kit (10-1247-01, Mercodia) according to the manufacturer’s instructions.

### Oral glucose tolerance test

An oral glucose tolerance test (OGTT) was performed in offspring at 7 weeks of age. Mice were fasted for 6 h and glucose was administered orally by gavage at 1 g/kg body weight. Blood glucose levels were measured at 0, 15, 30, 45, and 60 min using a laboratory animal glucose meter (Lab Gluco, RIJ). The incremental area under the curve (iAUC) relative to baseline glucose was calculated using the trapezoidal method.

An insulin tolerance test (ITT) was performed in offspring at 8 weeks of age, 1 week after the OGTT. Mice were fasted for 2 h and insulin was administered intraperitoneally at 0.75 U/kg body weight. Blood glucose levels were measured at 0, 15, 30, 45, and 60 min using the glucose meter. The area above the curve (AAC) relative to baseline glucose was calculated using the trapezoidal method.

### Pancreatic tissue preparation and histology

Pancreatic tissues were fixed in 10% neutral buffered formalin for 2 h. Samples were dehydrated sequentially in 90% ethanol for 2 h, 99.5% ethanol for 4 h, and 100% ethanol for 6 h, followed by clearing in xylene for 6 h and paraffin infiltration for 2 h. Tissue processing was performed using an automated paraffin-processing system (RH-12DM, Sakura Finetek Japan Co., Ltd., Japan). Paraffin-embedded tissues were sectioned at 4 μm using a microtome (SM2010R, Leica Biosystems, Japan). For hematoxylin and eosin (H&E) staining, sections were deparaffinized in xylene, washed under running water, stained with hematoxylin for 5 min and eosin-alcohol solution for 2 min, dehydrated through alcohol, cleared in xylene, and mounted. For immunohistochemical detection of insulin and glucagon, paraffin-embedded pancreatic sections were deparaffinized in xylene and incubated in 90% methanol containing 10% hydrogen peroxide for 4 min to suppress endogenous peroxidase activity. Sections were washed three times for 5 min each with TBS and blocked with skim milk at 37°C for 40 min. Sections were incubated overnight at 4°C with primary antibodies against insulin or glucagon. Antibody information, including manufacturer, catalog number, and dilution, is provided in Supplementary Table S1. Secondary antibody detection was performed for 1 h using an EnVision HRP-labeled polymer detection system (Agilent/Dako; Anti-Mouse, K4001; Anti-Rabbit, K4003). Immunoreactivity was visualized by incubation with a DAB solution containing 0.5 mg/mL DAB, 50 mM Tris-HCl (pH 7.6), and 0.033% H_2_O_2_ for 5–10 min. Sections were subsequently counterstained with Mayer’s hematoxylin for 2 min, dehydrated, cleared in xylene, and mounted. H&E- and immunohistochemically stained sections were examined using a bright-field microscope (LMDF, Leica).

### Skeletal muscle immunofluorescence and morphometric analysis

Frozen TA muscles were transversely sectioned at 8 μm using a cryostat. For laminin immunostaining, sections were fixed with 4% paraformaldehyde for 5 min, permeabilized with PBS containing 0.1% Triton X-100 for 5 min, and blocked with Blocking One (Nacalai Tesque, Japan) for 5 min. Sections were incubated with primary antibodies overnight at 4°C, followed by incubation with appropriate secondary antibodies for 1 h. Nuclei were counterstained with DAPI (1:5,000; Dojindo).

For myofiber-type analysis, sections were fixed in ice-cold acetone for 5 min and blocked with M.O.M. Mouse IgG Blocking Reagent (Vector Laboratories, Belgium) for 1 h. Sections were incubated with primary antibodies overnight at 4°C, washed three times with PBS for 5 min each, incubated with appropriate secondary antibodies for 1 h, and mounted using SlowFade Diamond Antifade Reagent (Thermo Fisher Scientific). Antibody information, including manufacturers, catalog numbers, and dilutions, is provided in Supplementary Table S1.

Fluorescence images were acquired using an A1Rsi confocal microscope (Nikon, Japan). Whole transverse sections of the TA muscle were imaged by tile scanning. Myofiber cross-sectional area (CSA), total myofiber number, and the number of type I MyHC-positive myofibers were quantified using WinROOF 2021 (Mitani Corporation, Tokyo, Japan). The proportion of type I MyHC-positive myofibers was calculated relative to the total number of myofibers.

### RNA extraction and quantitative PCR

Total RNA was extracted from liver and skeletal muscle tissues using the RNeasy Mini Kit (Qiagen, USA). Tissues were homogenized in 700 μL QIAzol Lysis Reagent, followed by the addition of 140 μL chloroform. Samples were centrifuged at 12,000 × g for 15 min at 4°C, and the aqueous phase was collected. After addition of 1.5 volumes of 100% ethanol, total RNA was purified using RNeasy Mini spin columns according to the manufacturer’s instructions. RNA concentration was measured using a NanoDrop 1000 spectrophotometer (Thermo Fisher Scientific).

For cDNA synthesis, 300 ng of total RNA was reverse-transcribed using ReverTra Ace reverse transcriptase (TOYOBO, Japan) and random primers. Reverse transcription was performed using a TP600 thermal cycler (Takara Bio, Japan) at 30°C for 10 min, 42°C for 60 min, and 99°C for 5 min.

Quantitative real-time PCR was performed using THUNDERBIRD Next SYBR Mix (QPX-201, TOYOBO) on a StepOnePlus Real-Time PCR System (Applied Biosystems, Thermo Fisher Scientific). Following initial denaturation at 95°C for 1 min, amplification was performed for 50 cycles of 95°C for 15 s and 60°C for 45 s. Relative mRNA expression levels were calculated using the 2^−ΔΔCt method, with *Gapdh* as the reference gene. Primer sequences are listed in Supplementary Table S2.

### RNA sequencing

Total RNA was extracted from tibialis anterior (TA) muscles using the RNeasy Micro Kit (74004, Qiagen). RNA sequencing library preparation and sequencing were performed by Takara Bio Inc. (Japan). cDNA was generated using a SMART (Switching Mechanism at the 5′ End of RNA Template)-based method, followed by PCR amplification and purification using AMPure XP beads (Beckman Coulter). Sequencing libraries were generated by adapter and index addition followed by PCR amplification. Libraries were sequenced using a NovaSeq 6000 system (Illumina).

### Differential gene expression analysis

Base calling was performed using bcl2fastq2 (v2.20, Illumina) from raw NovaSeq 6000 sequencing data generated with NovaSeq Control Software v1.7.5 and Real-Time Analysis v3.4.4. Sequencing reads were mapped to the mouse reference genome (GRCm39, GENCODE release M28), and gene-level read counts were generated using the DRAGEN Bio-IT Platform (v3.9.3, Illumina) with default parameters and GENCODE M28 gene annotation (Takara Bio Inc.). Gene-level counts were analyzed separately for the 4- and 8-week time points (n = 3 per group) using DESeq2 (v1.46.0) in R. Genes with fewer than 10 total counts across the six samples at each time point were excluded prior to testing. Differential expression between DMO and Control groups was assessed using the DESeq2 Wald test (design formula: ∼ group). Log2 fold changes were shrunk using the apeglm method (v1.28.0) to stabilize estimates for low-count genes. Genes with a Benjamini– Hochberg-adjusted P value (padj) < 0.05 were considered differentially expressed. Complete differential expression analysis results are provided in Supplementary Table S3.

### Pathway enrichment analysis

Over-representation analysis of KEGG and Reactome pathways was performed separately for up- and down-regulated differentially expressed genes (DEGs; DESeq2, padj < 0.05) at 4 and 8 weeks. Ensembl gene identifiers were mapped to Entrez Gene IDs using bitr (clusterProfiler v4.14.6). For each analysis, the background was restricted to genes passing DESeq2 independent filtering (non-missing adjusted p-values) at the corresponding timepoint.

KEGG pathway enrichment was performed using enrichKEGG (organism = “mmu”; clusterProfiler v4.14.6), and Reactome pathway enrichment was performed using enrichPathway (organism = “mouse”; ReactomePA v1.50.0, reactome.db v1.89.0). P values were adjusted using the Benjamini–Hochberg method, and gene-set sizes were restricted to 10–500 genes.

Main figures show up to 20 pathways with padj < 0.01. KEGG disease-related pathways driven predominantly by shared oxidative-phosphorylation, proteasome, or ribosome genes were excluded, and the 8-week down-regulated Reactome results were restricted to pathways related to mitochondrial respiration, translation, and biogenesis. Complete significant enrichment results (padj < 0.05) and unrestricted 8-week down-regulated KEGG and Reactome dot plots are provided in Supplementary Table S4 and Supplementary Figure 1, respectively.

Dot plots show fold enrichment, adjusted p-values, and gene counts and were generated using enrichplot::dotplot (v1.26.6) and ggplot2. All analyses were performed in R v4.4.2.

### DNA methylation analysis

Genome-wide DNA methylation was profiled in Control and DMO TA muscle at 4 weeks of age (n = 3/group) by reduced representation bisulfite sequencing (RRBS). Genomic DNA was extracted using the NucleoSpin Tissue kit (U0952S, Takara Bio). Library preparation and sequencing were performed by Rhelixa Inc. (Japan). DNA was digested with MspI, and libraries were prepared using the Zymo-Seq RRBS Library Kit (D5460/D5461, Zymo Research) and sequenced on a NovaSeq 6000 system (Illumina) with 150-bp paired-end reads.

Raw sequence reads were quality-checked using FastQC v0.11.7, trimmed using Trim Galore v0.5.0, and aligned to the mouse reference genome using Bismark v0.20.0. DNA methylation levels and differential methylation were analyzed using methylKit v1.16.0. For region-level analysis, the genome was divided into non-overlapping 1,000-bp windows. Differentially methylated regions (DMRs) and differentially methylated cytosines (DMCs) were defined by a q-value < 0.01 and an absolute methylation difference >25% between the Control and DMO groups. Each DMR/DMC was annotated to its nearest gene and genomic context (promoter, exon, intron, intron-exon boundary, 3′UTR, or intergenic).

For each significant 4-week DEG (padj < 0.05), associated methylation changes were classified as Tier 1 (region-level DMR) or Tier 2 (single-CpG DMC without a corresponding DMR). For genes with multiple DMCs, the site with the lowest q-value was used. Concordance between methylation and gene expression was defined as hypermethylation with decreased expression or hypomethylation with increased expression and was tested against an expected frequency of 50% using a two-sided exact binomial test (R v4.4.2).

### Statistical analysis

Data are presented as individual values with bars indicating the mean. Homogeneity of variance between two groups was assessed using an F-test. Comparisons between the Control and DMO groups were performed using an unpaired Student’s *t*-test, with Welch’s correction when variances differed between groups. For time-course experiments, data were analyzed by two-way ANOVA followed by Šídák’s multiple-comparisons test to compare Control and DMO groups at each time point. A *p* value < 0.05 was considered statistically significant. Statistical analyses were performed using GraphPad Prism version 10.3.1 (GraphPad Software).

## Results

### Maternal hyperglycemia alters postnatal growth and blood glucose levels in offspring

To investigate the effects of maternal hyperglycemia on postnatal growth and metabolism in offspring, hyperglycemia was induced in female mice by three consecutive intraperitoneal injections of streptozotocin (STZ; 100 mg/kg), and male offspring were analyzed at 4 and 8 weeks of age (Fig. 1A). Maternal body weight was comparable between the groups before STZ administration but was significantly lower in hyperglycemic mothers from gestational day 5 onward (Fig. 1B). Maternal blood glucose levels were significantly higher in the DMO group than in the control group from gestational day 5 onward (Fig. 1C). Litter size was also significantly smaller in hyperglycemic mothers than in control mothers (Fig. 1D).

**Figure 1.**
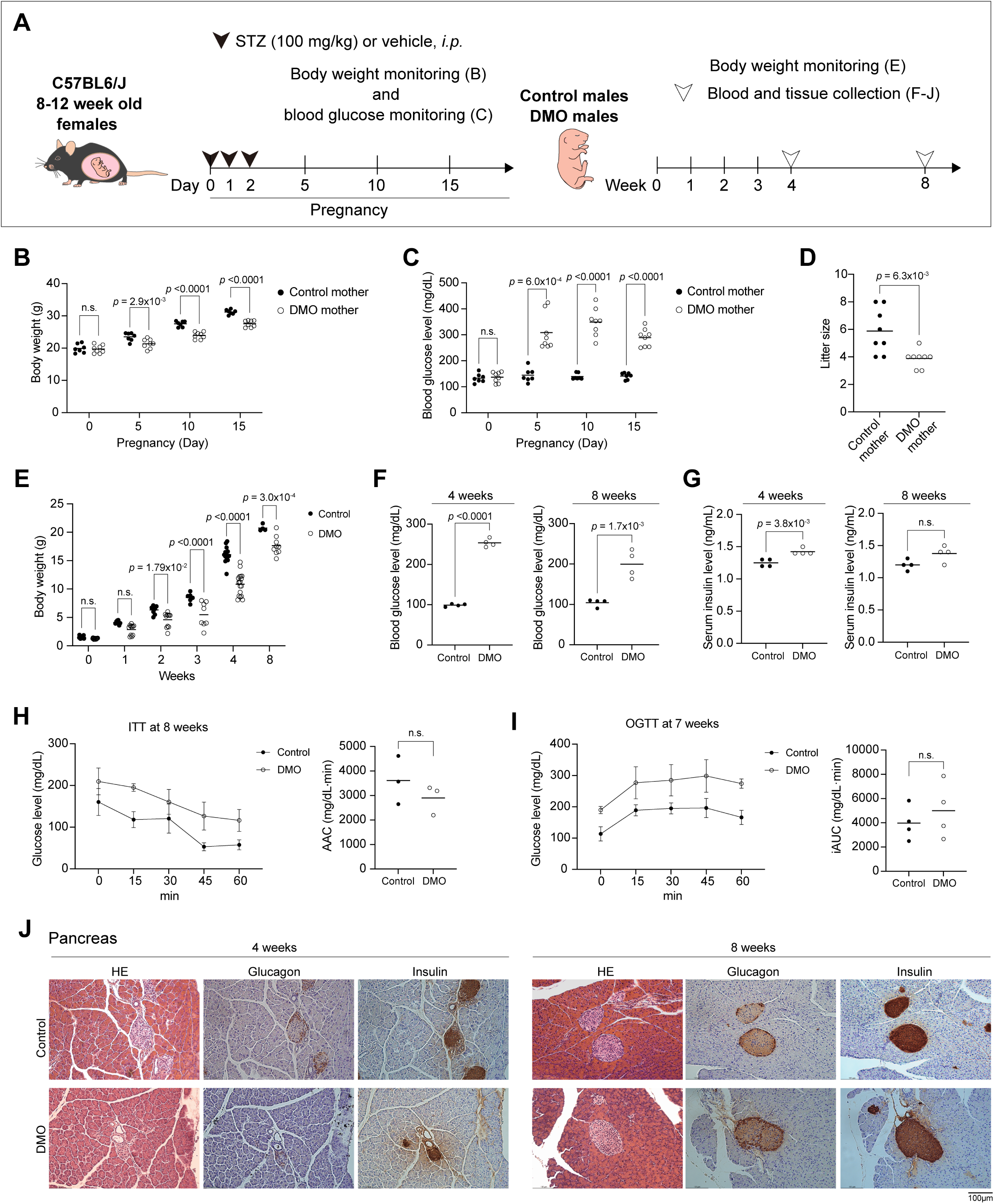
Maternal hyperglycemia alters postnatal growth and blood glucose levels in offspring. (A) Schematic overview of the experimental design. Female C57BL/6J mice aged 8–12 weeks received streptozotocin (STZ; 100 mg/kg, i.p.) or vehicle during early pregnancy. Maternal body weight and blood glucose levels were monitored during pregnancy. Male offspring born to vehicle-treated control mothers (Control) and hyperglycemic mothers following STZ treatment (DMO) were monitored after birth, subjected to metabolic tolerance tests at 7 and 8 weeks of age, and analyzed at 4 and 8 weeks of age. (B, C) Maternal body weight (B) and blood glucose levels (C) during pregnancy in Control and DMO mothers. (D) Litter size of Control and DMO mothers. (E) Body weight of male Control and DMO offspring from birth to 8 weeks of age. (F) Blood glucose levels in Control and DMO offspring at 4 and 8 weeks of age. (G) Serum insulin levels in Control and DMO offspring at 4 and 8 weeks of age. (H) Insulin tolerance test (ITT) in Control and DMO offspring at 8 weeks of age. Blood glucose levels were measured at 15-min intervals for 60 min following intraperitoneal insulin administration, and the area above the curve (AAC) relative to baseline glucose was calculated. (I) Oral glucose tolerance test (OGTT) in control and DMO offspring at 7 weeks of age. Blood glucose levels were measured at 15-min intervals for 60 min following oral glucose administration, and the incremental area under the curve (iAUC) relative to baseline glucose was calculated. (J) Representative pancreatic sections from Control and DMO offspring at 4 and 8 weeks of age stained with hematoxylin and eosin (H&E) or subjected to immunohistochemical staining for glucagon and insulin using DAB detection. Scale bar, 100 μm. Data are presented as mean ± SD. Each dot represents one individual animal. Comparisons between Control and DMO groups were performed using an unpaired Student’s t-test when variances were equal and Welch’s t-test when variances were unequal. For time-course data in B, C, and E, group differences at each time point were assessed using Šídák’s multiple-comparisons test following two-way ANOVA. Exact *p* values are shown in the figure. n.s., not significant.

Body weight did not significantly differ between control and DMO offspring at birth but was significantly lower in DMO offspring from 2 to 8 weeks of age (Fig. 1E). Blood glucose levels were significantly higher in DMO offspring than in controls at 4 and 8 weeks of age (Fig. 1F). Serum insulin levels were significantly higher in DMO offspring at 4 weeks, whereas no significant difference was observed at 8 weeks (Fig. 1G).

During the insulin tolerance test at 8 weeks, blood glucose remained higher in DMO offspring at several time points after insulin administration, but the area above the curve (AAC) did not significantly differ between groups (Fig. 1H). During the oral glucose tolerance test at 7 weeks, DMO offspring likewise had higher blood glucose throughout the test, whereas the incremental area under the curve (iAUC) did not significantly differ (Fig. 1I). Thus, DMO offspring were hyperglycemic during both tests, without a significant difference in glucose excursion. Histological examination of the pancreas and immunohistochemical staining for insulin and glucagon revealed no obvious morphological abnormalities in DMO offspring at 4 or 8 weeks of age (Fig. 1J). Offspring exposed to maternal hyperglycemia therefore showed reduced postnatal growth with elevated blood glucose levels.

### Hepatic alterations in DMO offspring are more pronounced at 4 weeks than at 8 weeks

Given the elevated blood glucose levels observed in DMO offspring, we first examined hepatic alterations associated with maternal hyperglycemia. Serum AST and ALT levels were significantly elevated in DMO offspring at 4 weeks of age, whereas no significant differences were observed at 8 weeks (Fig. 2A).

**Figure 2.**
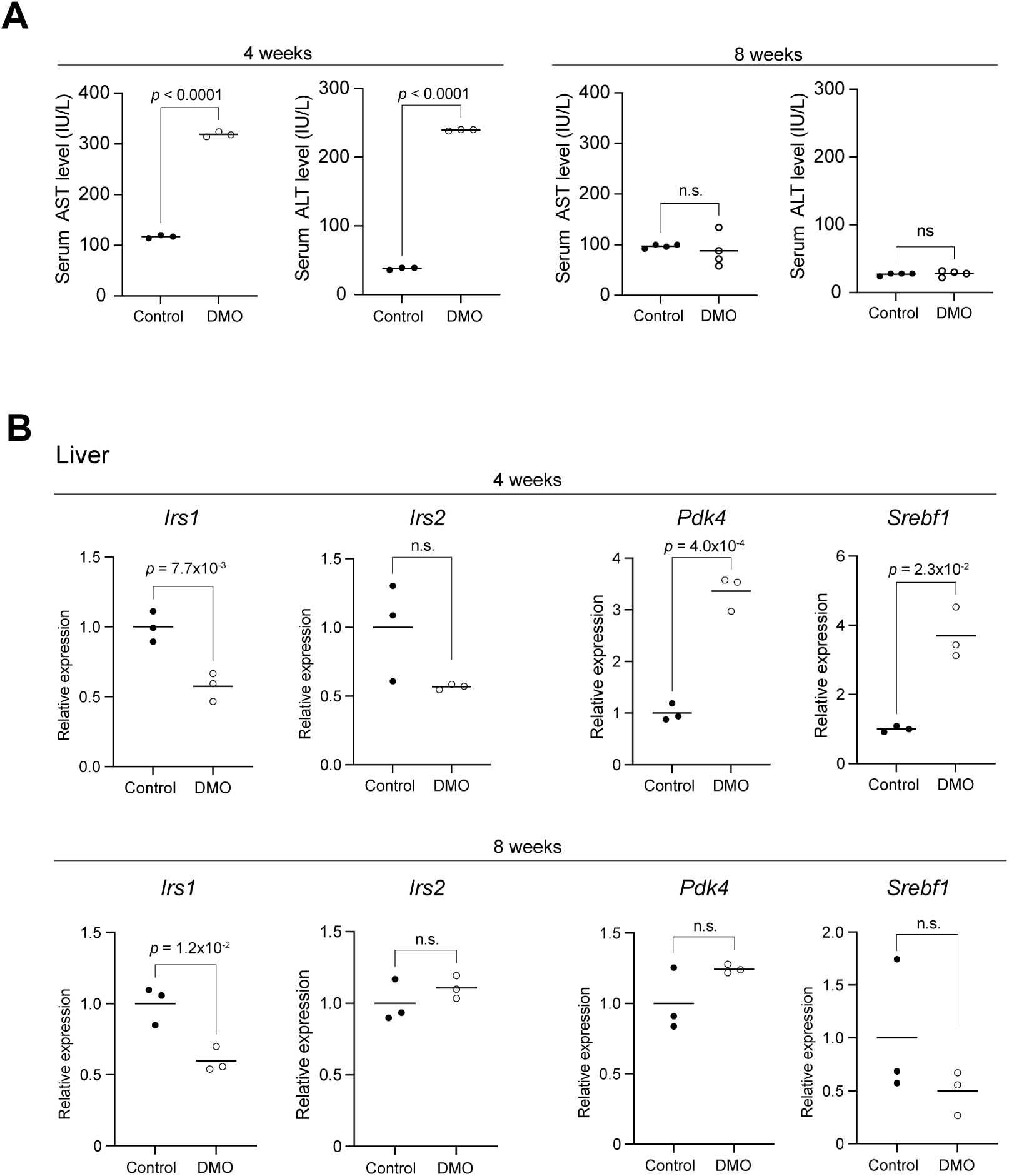
Postnatal stage-dependent hepatic alterations in offspring exposed to maternal hyperglycemia. (A) Serum aspartate aminotransferase (AST) and alanine aminotransferase (ALT) levels in Control and DMO offspring at 4 and 8 weeks of age. (B) Relative mRNA expression of *Irs1*, *Irs2*, *Pdk4*, and *Srebf1* in the liver of Control and DMO offspring at 4 and 8 weeks of age, determined by quantitative PCR. Data are presented as mean ± SD. Each dot represents one individual animal derived from a different litter. Comparisons between Control and DMO groups were performed using an unpaired Student’s t-test when variances were equal and Welch’s t-test when variances were unequal. Exact *p* values are shown in the figure. n.s., not significant.

We then examined the expression of genes involved in insulin signaling (*Irs1* and *Irs2*) and the regulation of glucose and lipid metabolism (*Pdk4* and *Srebf1*). At 4 weeks, *Irs1* expression was significantly decreased, whereas *Pdk4* and *Srebf1* expression was significantly increased in DMO offspring compared with controls. *Irs2* expression did not significantly differ between the groups (Fig. 2B). At 8 weeks, *Irs1* expression remained significantly lower in DMO offspring, whereas *Irs2*, *Pdk4*, and *Srebf1* expression did not significantly differ (Fig. 2B). Hepatic alterations in DMO offspring were therefore more evident at 4 weeks than at 8 weeks.

### Maternal hyperglycemia impairs postnatal skeletal muscle growth in offspring

We next focused on skeletal muscle and examined muscle weight and myofiber morphology to determine whether maternal hyperglycemia affected postnatal muscle growth. The weights of the gastrocnemius, tibialis anterior (TA), and soleus muscles were significantly lower in DMO offspring than in controls at 4 and 8 weeks of age (Fig. 3A). The combined length of the femur and tibia was significantly shorter in DMO offspring at 4 weeks, whereas no significant difference was observed at 8 weeks (Fig. 3B).

**Figure 3.**
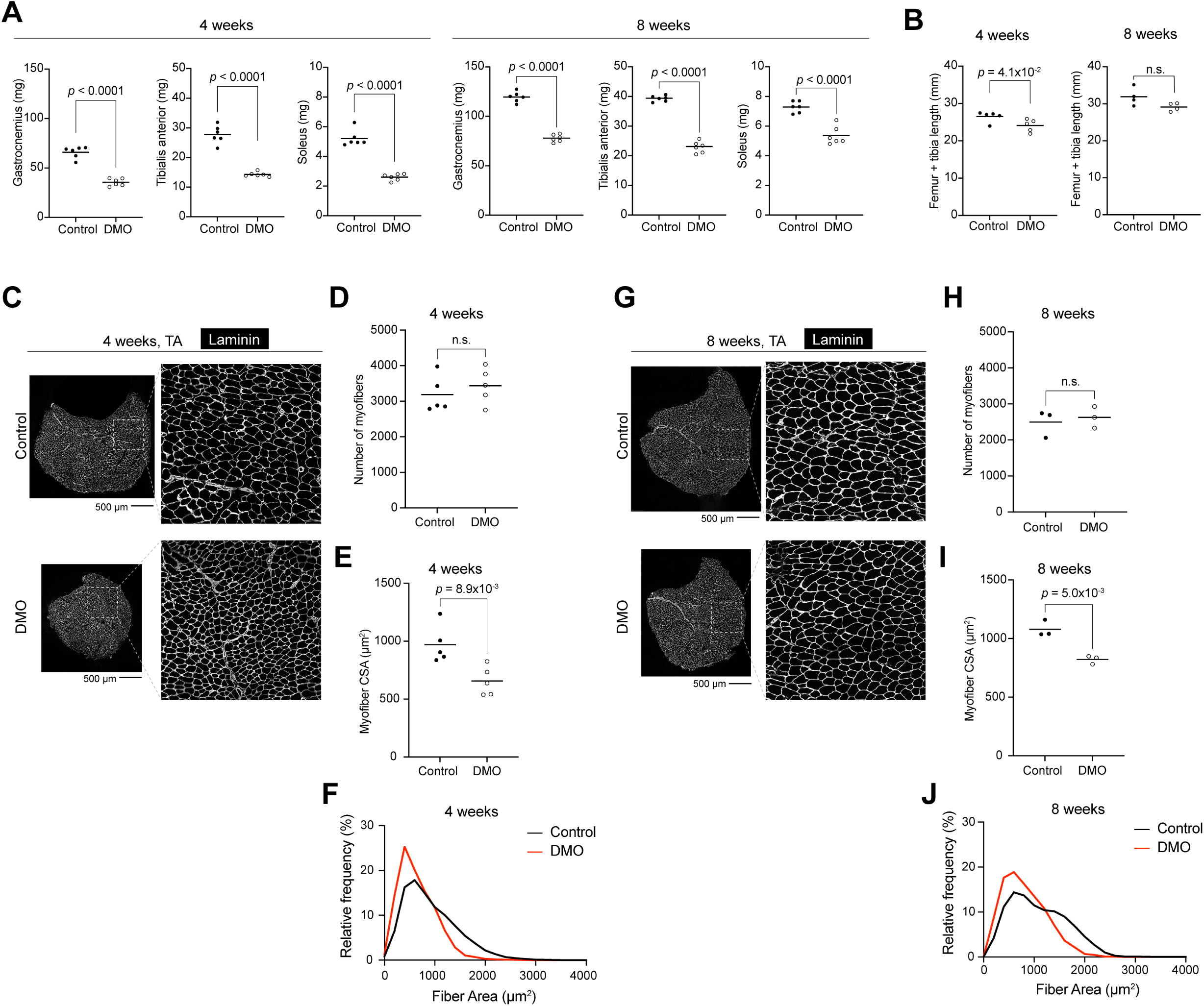
Maternal hyperglycemia impairs postnatal skeletal muscle growth in offspring. (A) Gastrocnemius, tibialis anterior (TA), and soleus muscle weights in Control and DMO offspring at 4 and 8 weeks of age. (B) Combined length of the femur and tibia in Control and DMO offspring at 4 and 8 weeks of age. (C, G) Representative transverse sections of TA muscles from Control and DMO offspring at 4 weeks (C) and 8 weeks (G) of age immunostained for laminin. Whole-muscle sections and higher-magnification images of the indicated regions are shown. Scale bars, 500 μm. (D, H) Total number of myofibers in TA muscle cross-sections at 4 weeks (D) and 8 weeks (H) of age. (E, I) Mean myofiber cross-sectional area (CSA) in TA muscles at 4 weeks (E) and 8 weeks (I) of age. (F, J) Distribution of myofiber CSA in TA muscles at 4 weeks (F) and 8 weeks (J) of age. Data are presented as mean ± SD. Each dot represents one individual animal derived from a different litter. Comparisons between Control and DMO groups were performed using an unpaired Student’s t-test when variances were equal and Welch’s t-test when variances were unequal. Exact *p* values are shown in the figure. n.s., not significant.

TA muscle morphology was examined by laminin immunostaining (Fig. 3C, G). The total number of myofibers in TA cross-sections did not significantly differ between control and DMO offspring at either time point (Fig. 3D, H). In contrast, mean myofiber cross-sectional area (CSA) was significantly smaller in DMO offspring at 4 and 8 weeks (Fig. 3E, I). Consistent with this finding, the distribution of myofiber CSA was shifted toward smaller fiber sizes in DMO offspring at both time points (Fig. 3F, J). Reduced skeletal muscle growth in DMO offspring was therefore associated with impaired postnatal myofiber hypertrophy rather than a reduction in myofiber number.

### Maternal hyperglycemia alters skeletal muscle fiber-type composition in offspring

Because alterations in skeletal muscle fiber-type composition have been associated with diabetes and metabolic dysfunction [16], we next examined myofiber-type composition in DMO offspring. We first analyzed the expression of myosin heavy chain genes in TA muscles. At 4 weeks, the expression levels of *Myh1*, *Myh2*, *Myh4*, and *Myh7* did not significantly differ between control and DMO offspring (Fig. 4A left). At 8 weeks, *Myh7* expression was significantly increased in DMO offspring, whereas no significant differences were observed in *Myh1*, *Myh2*, or *Myh4* expression (Fig. 4A right).

**Figure 4.**
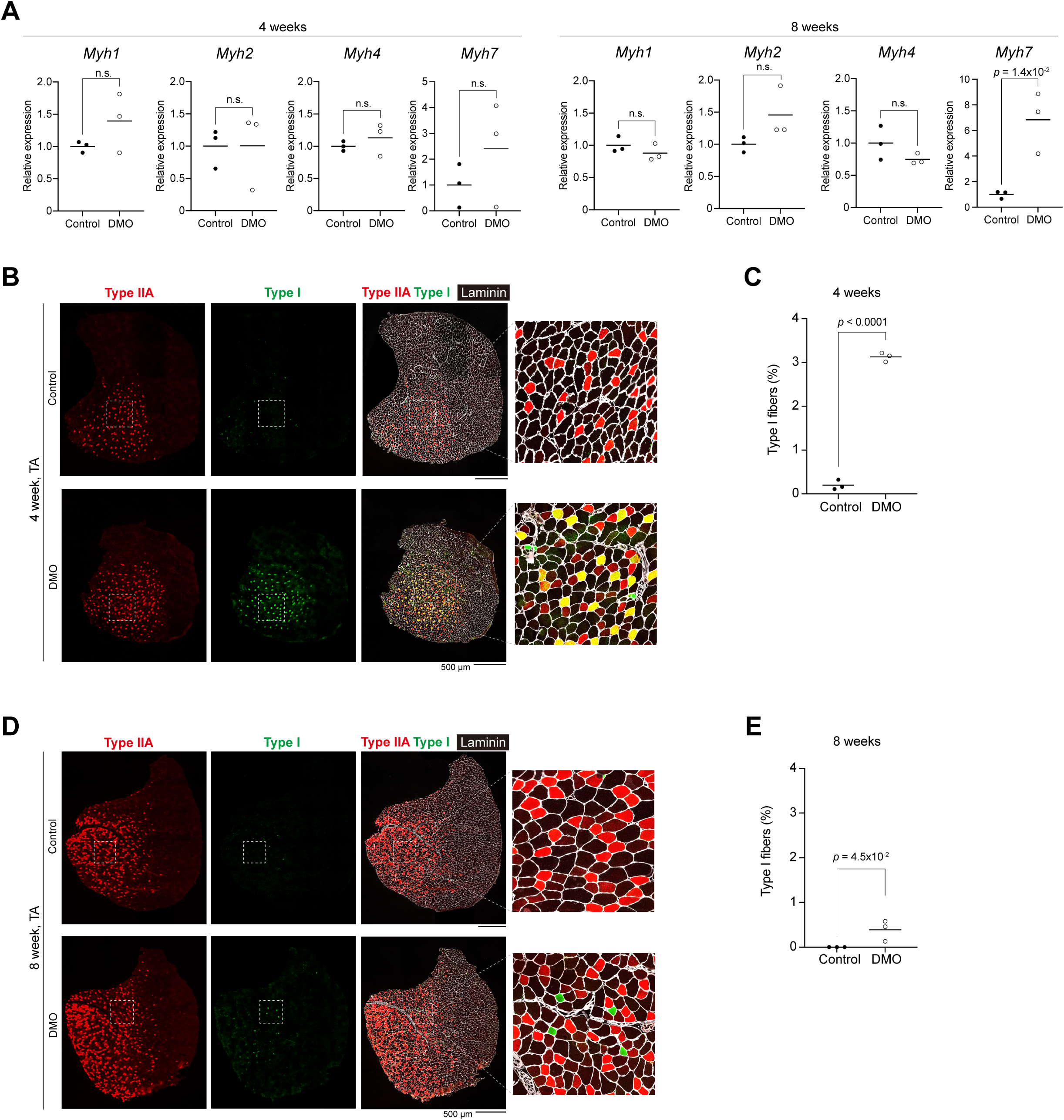
Maternal hyperglycemia alters skeletal muscle fiber-type composition in offspring. (A) Relative mRNA expression of *Myh1*, *Myh2*, *Myh4*, and *Myh7* in TA muscles from Control and DMO offspring at 4 and 8 weeks of age, determined by quantitative PCR. (B, D) Representative transverse sections of TA muscles from Control and DMO offspring at 4 weeks (B) and 8 weeks (D) of age immunostained for type IIA myosin heavy chain (red), type I myosin heavy chain (green), and laminin (white). Higher-magnification images of the indicated regions are shown on the right. Scale bars, 500 μm. (C, E) Percentage of type I myofibers in TA muscles from Control and DMO offspring at 4 weeks (C) and 8 weeks (E) of age. Data are presented as mean ± SD. Each dot represents one individual animal derived from a different litter. Comparisons between Control and DMO groups were performed using an unpaired Student’s t-test when variances were equal and Welch’s t-test when variances were unequal. Exact *p* values are shown in the figure. n.s., not significant.

Myofiber types were then examined by immunostaining for type I and type IIA myosin heavy chains (Fig. 4B, D). At 4 weeks, type I/IIA hybrid myofibers, positive for both type I and type IIA myosin heavy chains and appearing yellow in the merged images, were readily observed in DMO TA muscles but were rarely detected in controls (Fig. 4B). The proportion of type I-positive myofibers was significantly higher in DMO offspring than in controls (Fig. 4C). At 8 weeks, type I-positive/type IIA-negative myofibers were observed in DMO TA muscles but were rarely detected in controls (Fig. 4D), and the proportion of type I-positive myofibers remained significantly higher in DMO offspring (Fig. 4E). Thus, DMO offspring showed a higher proportion of type I-positive myofibers at both time points, consisting of type I/IIA hybrid fibers at 4 weeks and type I-positive/type IIA-negative fibers at 8 weeks.

### Muscle growth impairment in DMO offspring is not accompanied by induction of muscle atrophy-related genes

As shown in Fig. 3, DMO offspring exhibited reduced skeletal muscle weight and myofiber CSA at 4 and 8 weeks of age. We therefore examined whether the expression of genes associated with muscle atrophy and autophagy was altered in DMO TA muscles. At 4 weeks, the expression of *Foxo1*, *Trim63*, *Fbxo32*, and *Egln3* was significantly lower in DMO TA muscles than in controls, whereas *Foxo3* and *Bnip3* expression did not significantly differ between groups (Fig. 5A). At 8 weeks, *Foxo1* and *Bnip3* expression was significantly lower in DMO offspring, whereas *Foxo3*, *Trim63*, *Fbxo32*, and *Egln3* expression did not significantly differ (Fig. 5B). Thus, despite the reduced myofiber size in DMO offspring, the examined atrophy- and autophagy-related genes were not coordinately upregulated at either time point.

**Figure 5.**
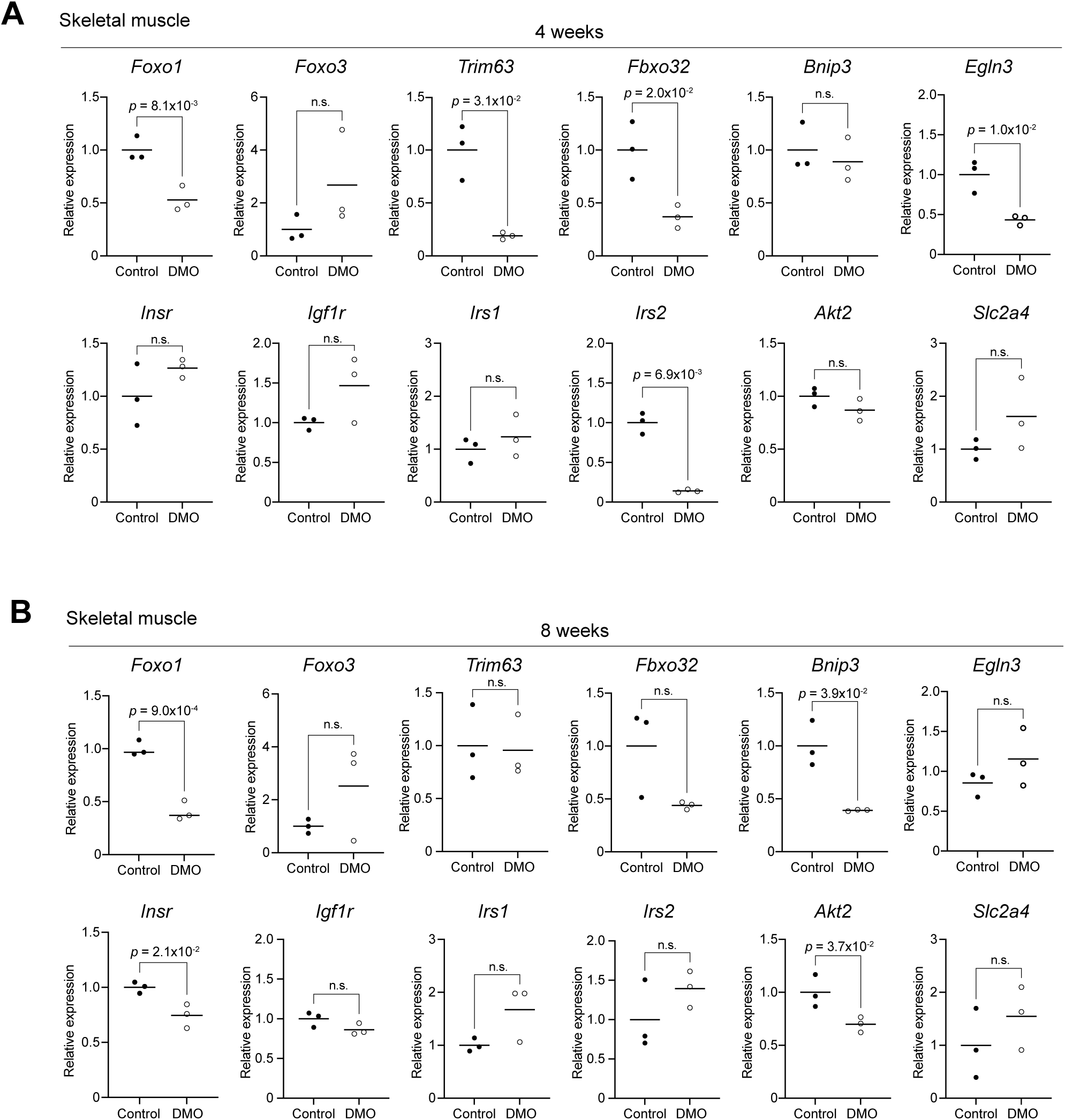
Alterations in insulin signaling and muscle atrophy-related gene expression in skeletal muscle of DMO offspring. (A, B) Relative mRNA expression of genes associated with muscle atrophy and autophagy (*Foxo1*, *Foxo3*, *Trim63*, *Fbxo32*, *Bnip3*, and *Egln3*) and genes involved in insulin/IGF signaling and glucose uptake (*Insr*, *Igf1r*, *Irs1*, *Irs2*, *Akt2*, and *Slc2a4*) in TA muscles from Control and DMO offspring at 4 weeks (A) and 8 weeks (B) of age, determined by quantitative PCR. Data are presented as mean ± SD. Each dot represents one individual animal derived from a different litter. Comparisons between Control and DMO groups were performed using an unpaired Student’s t-test when variances were equal and Welch’s t-test when variances were unequal. Exact *p* values are shown in the figure. n.s., not significant.

We also examined genes involved in insulin/IGF signaling and glucose uptake. At 4 weeks, *Irs2* expression was significantly decreased in DMO TA muscles, whereas *Insr*, *Igf1r*, *Irs1*, *Akt2*, and *Slc2a4* did not significantly differ between groups (Fig. 5A). At 8 weeks, *Insr* and *Akt2* expression was significantly decreased in DMO offspring, whereas *Igf1r*, *Irs1*, *Irs2*, and *Slc2a4* expression did not significantly differ (Fig. 5B). Impaired myofiber growth in DMO offspring therefore occurred without an overt atrophy-related transcriptional response and was accompanied by stage-dependent changes in insulin signaling-related gene expression.

### Transcriptomic analysis reveals postnatal stage-dependent alterations in DMO skeletal muscle

To further characterize the molecular alterations associated with impaired skeletal muscle growth in DMO offspring, we performed RNA sequencing of TA muscle at 4 and 8 weeks of age. PCA showed separation between control and DMO samples at both time points, with a more distinct separation at 4 weeks (Fig. 6A, C). Differential expression analysis identified 2,057 differentially expressed genes (DEGs) at 4 weeks, including 960 upregulated and 1,097 downregulated genes, and 644 DEGs at 8 weeks, including 278 upregulated and 366 downregulated genes (padj < 0.05; Fig. 6B, D).

**Figure 6.**
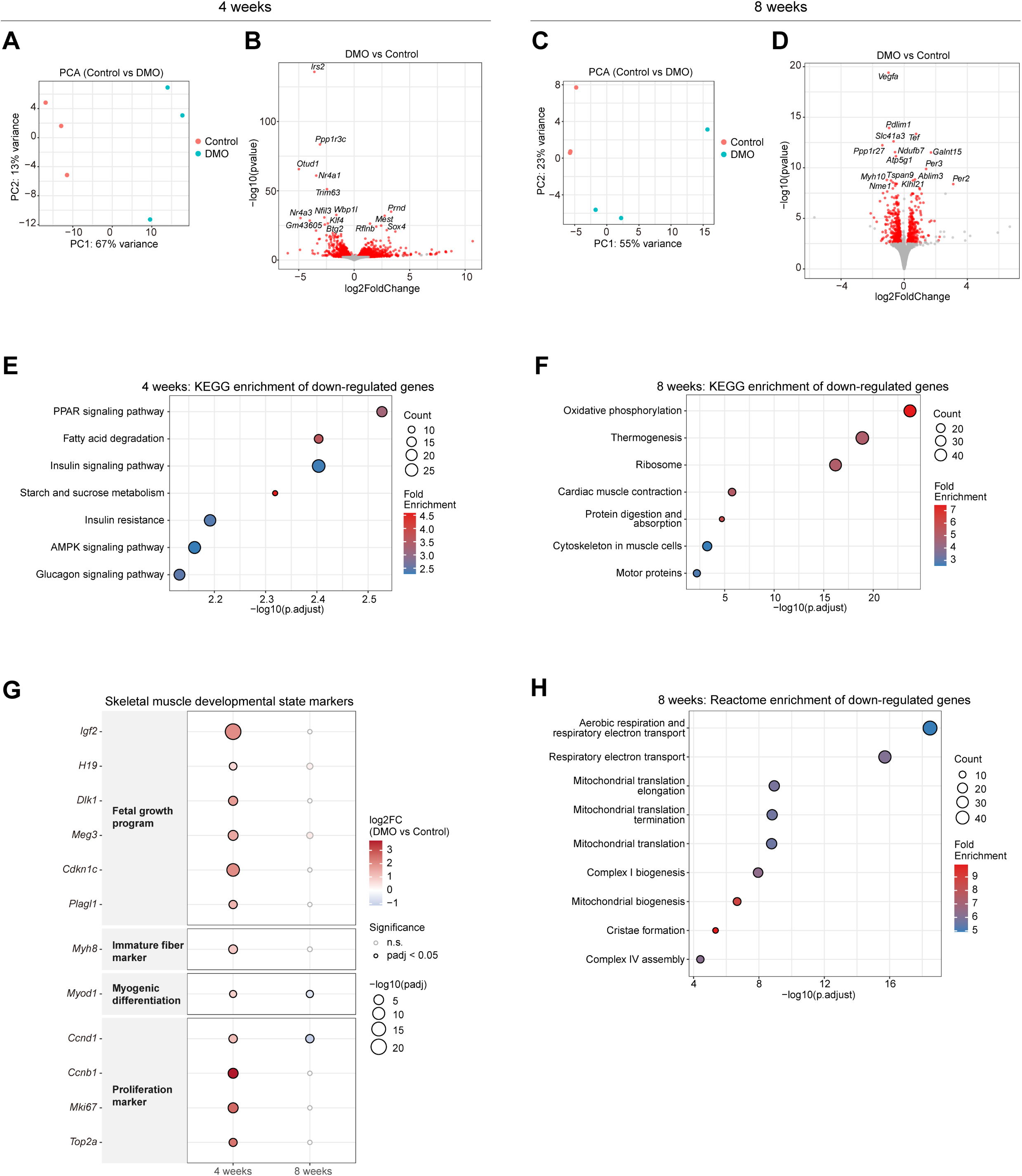
Transcriptomic alterations in DMO skeletal muscle at 4 and 8 weeks. (A, C) Principal component analysis (PCA) of variance-stabilized RNA-seq read counts from Control and DMO skeletal muscle at 4 weeks (A) and 8 weeks (C). n = 3 per group at each time point. (B, D) Volcano plots showing differential gene expression in DMO versus Control skeletal muscle at 4 weeks (B) and 8 weeks (D). The x-axis represents apeglm-shrunken log2 fold change, and the y-axis represents −log10(raw *P* value). Genes with an adjusted *P* value (padj) < 0.05 are shown in red; the 15 genes with the lowest padj at each time point are labeled. (E, F) KEGG pathway over-representation analysis of genes downregulated in DMO skeletal muscle (padj < 0.05) at 4 weeks (E) and 8 weeks (F). The background was restricted to genes passing DESeq2 independent filtering at the corresponding time point. Pathways are ranked by adjusted P value; dot color represents fold enrichment and dot size represents gene count. For the 8-week analysis (F), disease-related KEGG pathways driven predominantly by shared oxidative-phosphorylation, proteasome, or ribosome genes were excluded from the main figure for clarity. Complete KEGG enrichment results are provided in Supplementary Table S4. A complete dot plot of the 8-week downregulated KEGG results without pathway selection is shown in Supplementary Fig. S1A. (G) Expression changes in selected skeletal muscle developmental state markers at 4 and 8 weeks. Genes are grouped into fetal growth program (*Igf2, H19, Dlk1, Meg3, Cdkn1c,* and *Plagl1*), immature fiber (*Myh8*), myogenic differentiation (*Myod1*), and proliferation (*Ccnd1, Ccnb1, Mki67,* and *Top2a*) categories. Dot fill represents log2 fold change (DMO versus Control), dot size represents −log10(adjusted *P* value), and the outline indicates statistical significance (black, padj < 0.05; gray, not significant). (H) Reactome pathway over-representation analysis of genes downregulated in DMO skeletal muscle at 8 weeks, showing a selected subset of pathways related to mitochondrial respiration, translation, and mitochondrial biogenesis. Complete Reactome enrichment results are provided in Supplementary Table S4, and the complete dot plot without pathway selection is shown in Supplementary Fig. S1B. Dot color represents fold enrichment and dot size represents gene count.

Functional enrichment analyses revealed distinct patterns among upregulated and downregulated genes. At 4 weeks, upregulated genes were enriched for cell cycle-related pathways, including the KEGG cell cycle pathway (Supplementary Table S4). In contrast, downregulated genes showed broad enrichment for metabolic pathways. KEGG analysis identified PPAR signaling, fatty acid degradation, insulin signaling, AMPK signaling, and glucagon signaling (Fig. 6E; Supplementary Table S4).

Because reduced myofiber size and altered myofiber-type composition at 4 weeks could reflect changes in postnatal muscle development, we next examined selected genes associated with skeletal muscle developmental states. At 4 weeks, genes associated with the fetal growth program (*Igf2*, *H19*, *Dlk1*, *Meg3*, *Cdkn1c*, and *Plagl1*), immature myofibers (*Myh8*), myogenic differentiation (*Myod1*), and cell proliferation (*Ccnd1*, *Ccnb1*, *Mki67*, and *Top2a*) were significantly upregulated in DMO skeletal muscle (Fig. 6G). This coordinated upregulation was no longer observed at 8 weeks.

At 8 weeks, downregulated genes showed prominent enrichment for mitochondrial and oxidative processes. KEGG analysis identified oxidative phosphorylation and thermogenesis among the enriched pathways (Fig. 6F; Supplementary Table S4 and Supplementary Fig. S1A). Reactome analysis further identified pathways related to aerobic respiration and respiratory electron transport, mitochondrial translation, mitochondrial biogenesis, cristae formation, and respiratory chain complex assembly (Fig. 6H; Supplementary Table S4 and Supplementary Fig. S1B). Thus, DMO skeletal muscle showed distinct transcriptional alterations at the two time points, with developmental and metabolic changes at 4 weeks and predominant alterations in mitochondrial and oxidative metabolism at 8 weeks.

### DNA methylation changes explain only a limited proportion of transcriptional alterations at 4 weeks

Given the extensive transcriptional alterations observed in DMO skeletal muscle at 4 weeks, we next examined whether these changes were associated with alterations in DNA methylation. Among the 2,057 genes that were differentially expressed at 4 weeks, only 22 genes (1.07%) were associated with Tier 1 changes (region-level DMRs) (Fig. 7). An additional 436 genes (21.2%) were associated only with Tier 2 changes (single-CpG DMCs without a corresponding DMR), whereas 1,599 genes (77.73%) had no detectable methylation hit.

**Figure 7.**
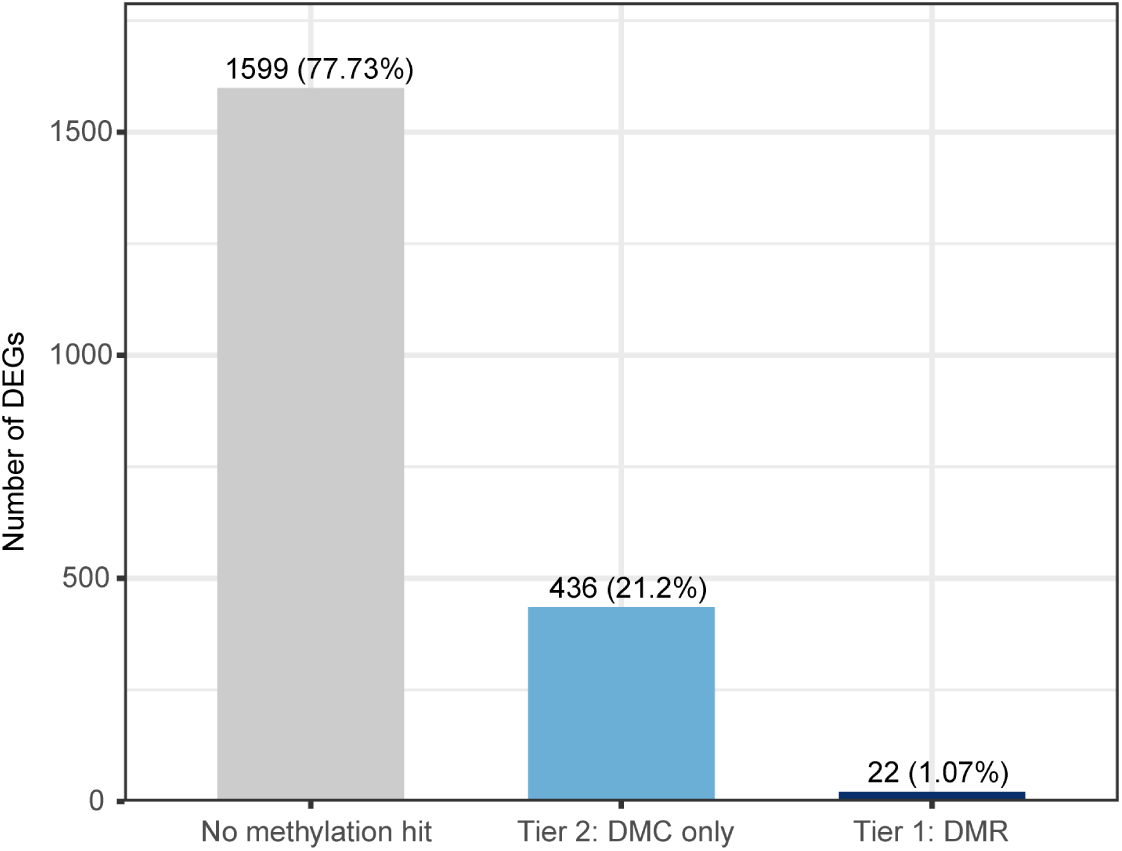
Coordinated 4-week DNA methylation changes are rare among 4-week DEGs. Distribution of 4-week-significant differentially expressed genes (DEGs; DESeq2, padj < 0.05; n = 2,057) according to their association with DNA methylation changes at 4 weeks. DEGs were classified as having no methylation hit, a Tier 2 hit (single-CpG DMC without a corresponding DMR), or a Tier 1 hit (region-level DMR). DMRs and DMCs were identified using a 25% methylation-difference threshold (see Methods). Gene-level details for Tier 1 and Tier 2 genes, together with a concordance analysis assessing whether the direction of methylation change was consistent with the expected inverse relationship with gene expression, are provided in Supplementary Table S5.

We further examined the directional relationship between DNA methylation and gene expression among the 458 genes associated with either Tier 1 or Tier 2 methylation changes. Of these genes, 53.9% (95% CI, 49.2–58.6%) showed concordance with the expected inverse relationship between DNA methylation and gene expression, defined as hypermethylation with decreased expression or hypomethylation with increased expression. This proportion did not significantly differ from 50% (binomial test, p = 0.10).

Coordinated DNA methylation changes at 4 weeks were therefore uncommon among differentially expressed genes and did not show a significant inverse relationship with gene expression. Gene-level annotations for Tier 1 and Tier 2 hits, and the concordance analysis, are provided in Supplementary Table S5.

## Discussion

Maternal hyperglycemia impaired postnatal skeletal muscle growth in offspring, with reduced muscle mass and myofiber size at both 4 and 8 weeks of age. The molecular phenotype, however, differed between stages. At 4 weeks, DMO muscle showed features associated with altered postnatal maturation, whereas mitochondrial and oxidative pathways were predominantly altered at 8 weeks. DNA methylation changes at 4 weeks showed only limited correspondence with differential gene expression. Thus, maternal hyperglycemia produced a persistent growth defect accompanied by stage-dependent molecular alterations.

The reduced postnatal growth and smaller litter size observed in DMO offspring are consistent with previous findings from severe experimental maternal diabetes. The direction of the growth phenotype in STZ-induced models varies with the timing and severity of maternal diabetes. Severe STZ-induced diabetes has been associated with fetal growth restriction, increased fetal loss, and smaller litters [3, 4, 6], while impaired postnatal growth has also been reported in offspring exposed to maternal diabetes [5]. Thus, the growth phenotype observed in the present study is consistent with the relatively severe maternal hyperglycemic environment produced in this model.

The smaller myofibers in DMO offspring were not accompanied by a reduction in fiber number, nor by coordinated induction of the examined atrophy- or autophagy-related genes [17, 18]. The growth phenotype therefore did not show the molecular features typically associated with a conventional atrophy response examined here. At 4 weeks, DMO muscle showed hybrid type I/IIA fibers and higher expression of fetal growth- and myogenesis-associated genes [19, 20], consistent with altered postnatal maturation. By 8 weeks, that signature was largely absent, and downregulated genes were enriched for mitochondrial and oxidative metabolism. Reduced expression of oxidative metabolism-related genes has also been reported in insulin-resistant and diabetic muscle [21, 22], and altered skeletal muscle transcriptional and metabolic profiles have also been reported in offspring exposed to intrauterine hyperglycemia [23]. Because DMO offspring remained hyperglycemic at 8 weeks, the later profile may reflect the ongoing metabolic environment as well as the earlier developmental effect. Persistent impairment of muscle growth was therefore not linked to a single molecular state.

Hepatic changes in this model were also more evident at 4 weeks than at 8 weeks, including elevated serum AST and ALT and altered expression of metabolic genes. Maternal hyperglycemia can affect the offspring liver [24–26], but here the hepatic phenotype was less persistent than the muscle growth defect.

Epigenetic modification, including DNA methylation, has been proposed as a mediator of lasting effects of maternal diabetes on offspring tissues [13–15]. Here, region-level methylation changes tagged only a small fraction of 4-week DEGs, and methylation changes did not show a significant inverse relationship with gene expression. Because both analyses were performed on whole skeletal muscle, however, cell-type-specific relationships may have been obscured. Thus, DNA methylation changes detectable by bulk analysis provided only a limited explanation for the early transcriptional phenotype.

In conclusion, severe maternal hyperglycemia in this STZ-induced model reduced skeletal muscle mass and myofiber size in male offspring at both postnatal stages examined. The accompanying molecular phenotype changed from a maturation-associated pattern at 4 weeks to reduced oxidative metabolism at 8 weeks. Concurrent DNA methylation showed limited correspondence with the 4-week transcriptional changes.

## Limitations

Several limitations should be considered. First, only male offspring were analyzed, and whether similar skeletal muscle phenotypes occur in females remains unknown. Second, diabetes was induced with STZ during early pregnancy. This model produces insulin-deficient maternal hyperglycemia and differs from the obesity-associated insulin resistance that commonly characterizes human gestational diabetes mellitus. Moreover, the maternal metabolic disturbance in the present model was relatively severe. Therefore, the reduced postnatal growth and skeletal muscle phenotypes observed here should not be directly extrapolated to human gestational diabetes or to milder maternal hyperglycemia. Third, fetal growth, fetal loss, and placental function were not measured, so the prenatal basis of the smaller litter size and reduced postnatal growth remains uncertain. Fourth, offspring remained with their biological dams until weaning. Prenatal hyperglycemia cannot be separated from lactational effects, including possible changes in milk yield or milk composition [27]. Fifth, RNA-seq and RRBS were performed on whole skeletal muscle, so cell-type-specific molecular changes cannot be distinguished from shifts in cellular composition. Sixth, DNA methylation was assessed only at 4 weeks, precluding analysis of earlier transient or later-emerging methylation changes.

## Abbreviations

AAC: area above the curve
CSA: cross-sectional area
DEG: differentially expressed gene
DMC: differentially methylated cytosine
DMO: diabetes mellitus offspring
DMR: differentially methylated region
iAUC: incremental area under the curve
ITT: insulin tolerance test
KEGG: Kyoto Encyclopedia of Genes and Genomes
OGTT: oral glucose tolerance test
PCA: principal component analysis
RNA-seq: RNA sequencing
RRBS: reduced representation bisulfite sequencing
STZ: streptozotocin
TA: tibialis anterior

## Author contributions

YT and TK conceived and designed the research, performed the experiments, analyzed the data, prepared the figures, and drafted the manuscript. YT, TK, TM, and MH interpreted the results of the experiments. DD provided advice on the transcriptomic analyses and contributed to interpretation of the results. YT, TK, MH, and DD edited and revised the manuscript. All authors approved the final version of the manuscript.

## Ethics Approval and Consent to Participate

All animal experiments were approved by the Animal Care and Use Committee of the Graduate School of Agricultural and Life Sciences, The University of Tokyo (approval no. P-21-107), and were performed in accordance with institutional guidelines for animal experimentation.

## Acknowledgments

We thank the Imaging Core Laboratory of the Institute of Medical Science, the University of Tokyo, for supporting this study. We also thank the InforBio Bioinformatics Platform at the Institut de Biologie Paris Seine, Sorbonne Université, for providing bioinformatics training that supported the data analysis performed in this study.

## Data Availability Statement

Sequencing data have been deposited in the database DDBJ at https://www.ddbj.nig.ac.jp under BioProject accession numbers PRJDB42664, PRJDB45732, and PRJDB45738.

## Supplemental Material

Supplementary Figure S1 and Supplementary Tables S1–S5 are provided as supplemental material. Supplementary Tables S3–S5 are provided as separate Excel files.

