## Supplemental Information for "Maternal hyperglycemia impairs postnatal skeletal muscle growth and induces stage-dependent gene expression changes in mouse offspring"

*Yayoi Tokunaga et al.*

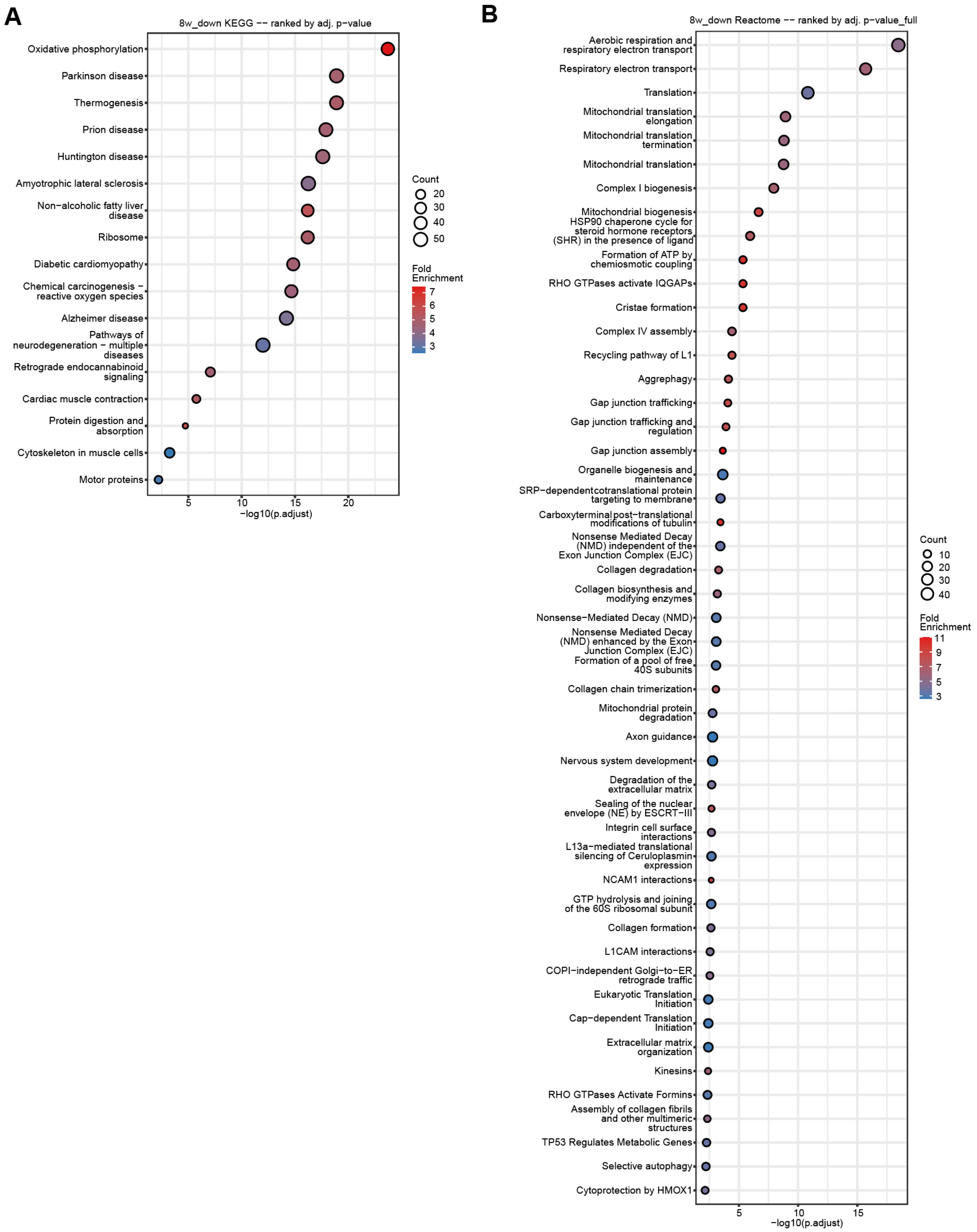


**Supplementary Figure S1. Unrestricted pathway enrichment results for 8-week down-regulated genes.**

Dot plots showing the complete significant pathway enrichment results (adjusted P value < 0.05) for 8-week down-regulated genes in (A) KEGG and (B) Reactome. Pathways are ranked by adjusted P value. Dot size indicates gene count, and color indicates fold enrichment.

**Supplementary Table S1. Antibodies used for immunohistochemistry and immunofluorescence.**

| Antibody | Host/isotype | Clone | Dilution | Manufacturer/ Database | Identifier |
|---|---|---|---|---|---|
| anti-insulin | Guinea pig | - | 1:200 | Dako | A0564 |
| anti-glucagon | Rabbit | - | 1:75 | Dako | A0565 |
| anti-MyHC I | Mouse IgG2b | BA-F8 | 1:4 | DSHB | ACC 154 |
| anti-MyHC IIA | Mouse IgG1 | SC-71 | 1:4 | DSHB | ACC 217 |
| anti-laminin | Rabbit IgG, polyclonal | - | 1:400 | Sigma-Aldrich | L9393 |
| Dylight 405 anti-mouse IgG2b | Goat IgG, polyclonal | - | 1:1000 | Jackson Immuno Research | 115-475-207 |
| Alexa Fluor® 555 anti-mouse IgG1 | Goat IgG | - | 1:1000 | Thermo Fisher Scientific | A21127 |
| Alexa Fluor® 488 anti-rabbit IgG | Donkey IgG, polyclonal | - | 1:1000 | Thermo Fisher Scientific | A32790 |

**Supplementary Table S2. Primer sequences used for quantitative PCR.**

| Gene | Direction | Sequence |
|---|---|---|
| *Gapdh* | Forward | 5’-CCTGGAGAAACCTGCCAAGT-3’ |
| | Reverse | 5’-AGAGTGGGAGTTGCTGTTGAAGTC-3’ |
| *Myh1* | Forward | 5’-GGAGGACCAAGTGAGTGAGC-3’ |
| | Reverse | 5’-TCGTCTAGCTGGCGTGAGTA-3’ |
| *Myh2* | Forward | 5’-AGCGACTGATCAACGACCTG-3’ |
| | Reverse | 5’-AACTGAGATACCAGCGCTTCC-3’ |
| *Myh4* | Forward | 5’-AAACCACCTCAGAGTTGTGGA-3’ |
| | Reverse | 5’-GTTCCGAAGGTTCCTGATTGC-3’ |
| *Myh7* | Forward | 5’-GCCCTTTGACCTCAAGAAAG-3’ |
| | Reverse | 5’-CTTCACAGTCACCGTCTTG-3’ |
| *Irs2* | Forward | 5’-ACTTCCCAGGGTCCCACTGCTG-3’ |
| | Reverse | 5’-GGCTTTGGAGGTGCCACGATAG-3’ |
| *Pdk4* | Forward | 5’-TACACGTACTCCACTGCTCC-3’ |
| | Reverse | 5’-CCTGACATAGAGTAGAGATT-3’ |
| *Srebf1* | Forward | 5’-AGCAGTCACCAGCTTCAGTC-3’ |
| | Reverse | 5’-GGTCATGTTGGAAACCACGC-3’ |
| *Irs1* | Forward | 5’-GAGGACTTGAGCTATGACAC-3’ |
| | Reverse | 5’-ATCAGGTTCTTTGTCTGACC-3’ |
| *Insr* | Forward | 5’-AAATGCAGGAACTCTCGGAAGCCT-3’ |
| | Reverse | 5’-ACCTTCGAGGATTTGGCAGACCTT-3’ |
| *Foxo1* | Forward | 5’-TGCTGTGAAGGGACAGATTG-3’ |
| | Reverse | 5’-GAGTGGATGGTGAAGAGCGT-3’ |
| *Foxo3* | Forward | 5’-ACAAACGGCTCACTTTGTCCCAGA-3’ |
| | Reverse | 5’-TCTTGCCCGTGCCTTCATTCT-3’ |
| *Trim63* | Forward | 5’-TGGAAACGCTATGGAGAACC-3’ |
| | Reverse | 5’-AAGATGTCGTTGGCACACTTC-3’ |
| *Fbxo32* | Forward | 5’-ACTTCTCGACTGCCATCCTG-3’ |
| | Reverse | 5’-GATGTGAGCTGTGACTTTGCTATC-3’ |
| *Bnip3* | Forward | 5’-CCCAGACACCACAAGATACCAACA-3’ |
| | Reverse | 5’-GGTCGACTTGACCAATCCCATATCCA-3’ |
| *Egln3* | Forward | 5’-ACGTGGAGCCCATTTTTGAC-3’ |
| | Reverse | 5’-CATAGCGTACCTGGTGGCATA-3’ |
| *Igf1r* | Forward | 5’-ATCGCGATTTCTGCGCCAACA-3’ |
| | Reverse | 5’-TTCTTCTCTTCATCGCCGCAGACT-3’ |
| *Akt2* | Forward | 5’-GCGCAAGGAGGTCATCATT-3’ |
| | Reverse | 5’-GCATACTTGAGGGCTGTAAGG-3’ |
| *Slc2a4* | Forward | 5’-CTTAGTAGAACGAGCTGGACGA-3’ |
| | Reverse | 5’-GACATAGCTCATGGCTGGAAC-3’ |

**Supplementary Table S3. Complete differential gene expression analysis results obtained using DESeq2.**

This table provides the complete results of differential gene expression analysis performed using DESeq2 for skeletal muscle samples at 4 and 8 weeks of age. Results for each time point are provided in separate worksheets. The table includes gene identifiers and annotations, baseMean, apeglm-shrunken log2 fold changes and their posterior standard errors (lfcSE), unshrunken log2 fold changes, Wald test P values, Benjamini–Hochberg-adjusted P values, and genomic coordinates. The complete results are provided separately as an Excel file.

**Supplementary Table S4. Complete KEGG and Reactome pathway enrichment results for differentially expressed genes.**

This table provides the complete significant KEGG and Reactome pathway enrichment results (adjusted P value < 0.05) for differentially expressed genes at 4 and 8 weeks of age. Results are provided in separate worksheets according to time point (4 or 8 weeks), direction of differential expression (up- or down-regulated), and pathway database (KEGG or Reactome). Pathways are ranked in ascending order of adjusted P values. The complete results are provided separately as an Excel file.

**Supplementary Table S5. Gene-level associations between differential gene expression and DNA methylation changes at 4 weeks.**

This table provides gene-level details of 4-week differentially expressed genes associated with DNA methylation changes. Tier 1 genes were associated with region-level differentially methylated regions (DMRs), whereas Tier 2 genes were associated with single differentially methylated cytosines (DMCs) only. Results for Tier 1 and Tier 2 genes are provided in separate worksheets. A concordance summary assessing whether the direction of DNA methylation change was consistent with the expected inverse relationship with gene expression is also provided. The complete results are provided separately as an Excel file.
